# Genomic fidelity and evolution of cancer patient-derived organoids

**DOI:** 10.64898/2025.12.10.693393

**Authors:** Linoy Raz, Haia Khoury, Tal Ben-Yishay, Ron Saad, Amjad Zoabi, Giulia Orlandi, Sara Donzelli, Giovanni Blandino, Uri Ben-David

**Affiliations:** Department of Human Molecular Genetics and Biochemistry, Faculty of Medicine, Tel Aviv University, Tel Aviv, Israel; The Blavatnik School of Computer Science, Faculty of Exact Sciences, Tel Aviv University, Tel Aviv, Israel; Translational Oncology Research Unit, IRCCS Regina Elena National Cancer Institute, Rome, 00144, Italy

## Abstract

**Background:** Patient-derived organoids (PDOs) are gaining recognition as a promising *ex vivo* model for cancer research, offering advantages over traditional 2D cell lines by better recapitulating tumor biology.

**Results:** In this study, we assess the genomic stability and evolution of PDOs by analyzing copy-number alterations (CNAs) in 300 PDO samples across 16 cancer types. These results are compared with data from previously analyzed patient-derived xenografts (PDXs). We observe that PDOs exhibit genomic evolution over passaging, with an increasing divergence from the original tumor genome over time. Importantly, across cancer types, PDOs maintain higher genomic fidelity and are more genetically similar to their tumors of origin than PDX models. Moreover, PDOs show greater genomic stability during culture passaging compared to PDXs.

**Conclusions:** These findings position PDOs as a reliable and representative model for cancer research, while highlighting the need to carefully track their genomic evolution in culture.

## Background

Patient-derived cancer models can evolve *in vitro* and *in vivo*, leading to genomic diversification of potential phenotypic consequence[1]. We and others have shown that cancer cell lines and patient-derived xenografts (PDXs) are subjected to distinct selection pressures that lead to their genomic evolution throughout their propagation[2– 6]. This genomic evolution can jeopardize the use of cancer models as ‘tumor avatars’[7–9], as well as the reproducibility of cancer research[10,11], with ramifications for the clinical translation of cancer research findings[12,13]. Therefore, characterizing the genetic stability of cancer models, understanding the extent to which they represent the original tumors from which they are derived, and documenting the extent of their genomic evolution throughout propagation, are key for the proper use of cancer models in biomedical research.

In recent years, patient-derived organoids (PDOs) have emerged as a widely used research model[14–16]. As three-dimensional (3D) *ex vivo* models of human tumors, PDOs were shown to better recapitulate various aspects of tumor biology in comparison to 2D models, including the response to mechanical forces, nutrient distribution, chromosome segregation[17], and even immune infiltration and vascularization[18–21]. However, the genomic resemblance of PDOs to their tumors of origin, as well as the degree of their genomic evolution throughout culture propagation, have not been comprehensively assessed across studies and tumor types. Further, how this genomic stability compares to that of PDXs – human tumors that grow in the mouse environment – has not been systematically studied. In this work, we performed a cross-cohort analysis of PDOs from 16 cancer types, comparing the landscape of copy number alterations (CNAs) between primary tumors and their derived PDOs, as well as between PDOs at different stages of their propagation (passage numbers). We also compared the genomic evolution of PDOs to that of PDXs, both by comparing matched PDOs and PDXs originating from the same primary tumors (PTs), and by comparing our PDO cohort to a large cohort of PDXs that we have previously characterized[2].

## Results

To characterize the genomic stability of PDOs, we collected whole-exome sequencing (WES) data from 300 PDO samples derived from 219 tumors of 205 individual patients across 13 studies of 16 cancer types[22–32]. This cohort includes 13 new PDOs that we derived from 11 patients and profiled using WES. Distribution of individual patients (models) and organoid samples by cancer type is shown in **Fig. 1A** and **Supplementary Fig. 1A**, respectively. Bladder, colorectal and gastric organoids are the most represented tumor types in our meta-cohort, which consists of 300 PT-PDO pairs, 23 of which also have a matched PDX and 17 of which have matched normal tissue samples (**Supplementary Fig. 1B**). Passage numbers were reported for ∼1/3 of the PDO models (**Supplementary Table 1**), and ranged from 0 to 31, with passage 8 being the median passage number (**Fig. 1B**).

**Fig 1.**
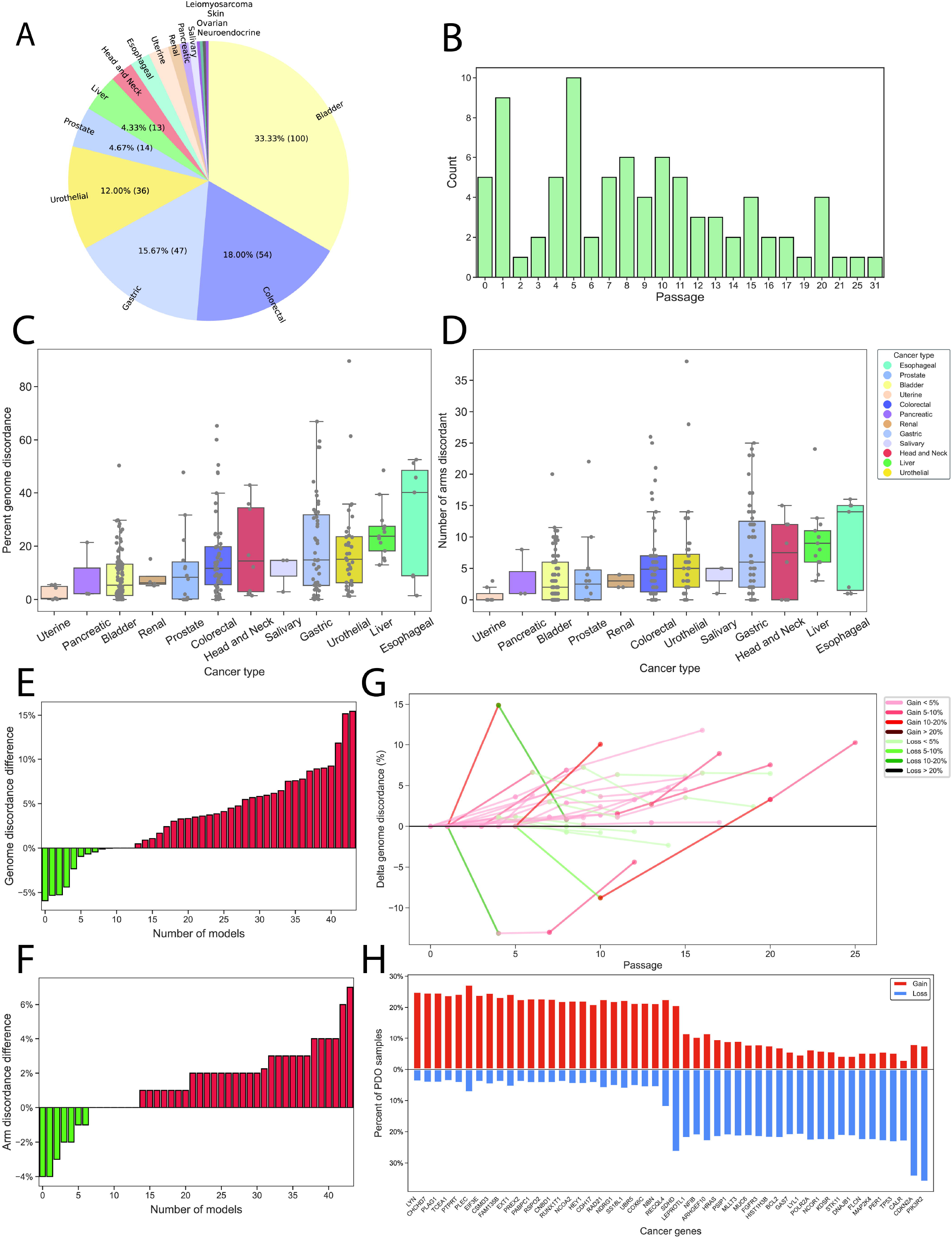
Characterization of copy-number discordance between PTs and PDOs. **(A)** Cancer type composition of the PDO samples included in this study. **(B)** Distribution of PDO passages across samples. **(C**,**D)** Percent of the genome that is discordant **(C)**, or the number of chromosome-arm CNAs that are discordant **(D)**, between PDOs and their PTs of origin, grouped by cancer type, in cohorts with at least 3 samples. A median of 13.05% of the genome was differentially altered between PDOs and PTs (median PDO of the median cohort). Kruskal-Wallis test; genome-level, p-value = 3.75e-07; Chromosome-arm level, p-value = 2.64e-06. Bar, median; colored rectangle, 25th to 75th percentile; whiskers, Q1–1.5*IQR to Q3+1.5*IQR. **(E**,**F)** The difference in the percent of the genome that is discordant **(E)** or the number of chromosome-arm CNAs that are discordant **(F)** between the earliest and latest matched PDO samples derived from the same tumor. Organoids in later passages are less similar to their tumors of origin than organoids in earlier passages, both at the bin-level and at the chromosome-arm-level. One-sided one-sample t-test; bin-level, p-value = 1.06e-05; Chromosome-arm level, p-value = 1.69e-04. **(G)** The cumulative PT-PDO genome discordance in a longitudinal series of matched PDOs. Each segment is colored by the change in discordance between the two consecutive organoids. The fraction of the genome that is discordant between passages is usually smaller than 5%, but there is an overall increase in the PT-PDO discordance throughout passaging. One-side one-sample t-test, p-value = 0.0002. **(H)** The top 25 differentially gained and lost cancer genes in PDO models. Cancer genes are significantly altered in PDOs - oncogenes are significantly gained while tumor suppressor genes are significantly lost. One-tailed Fisher’s exact test; Oncogene gain, p-value = 0.03; Tumor-suppressor gene loss, p-value < 0.0001.

We first derived copy number profiles from the WES data, using the established tool CNVkit[33] (**Methods**). Next, we compared the CN landscapes of PDOs to those of the PTs from which they were derived. Applying the same approach that we previously applied to PDXs[2,6], we inferred CNAs – both at a 1Mb resolution and at a chromosome-arm resolution – for each sample based on its WES data, and calculated the fraction of the genome that is differentially altered (FGA) by CNAs between each pair of samples, using conservative discordance thresholds (**Methods**). In our PDO cohort, we observed a median CNA difference of ∼13% of the genome (**Fig. 1C** and **Supplementary Fig. 2A**), or ∼5 chromosome-arms (**Fig. 1D and Supplementary Fig. 2B**), between the PDOs and their tumors of origin, across cancer types. The median discordance between PDOs and PTs was much higher for some cancer types (e.g., a median of ∼40% vs. ∼2% CN discordance for esophageal and pancreatic cancers, respectively; **Fig. 1C**), but this variability seems to be related mostly to the specific study rather than to the cancer type *per se*, as the PT-PDOs discordance varied greatly across studies within the same cancer type (see, for example, colorectal and bladder tumors; **Supplementary Fig. 2C,D**).

Next, we assessed the genomic evolution of PDOs throughout passaging, by comparing the fraction of the genome discordant (**Fig. 1E**) and the number of chromosome-arms discordant (**Fig. 1F**) between matched PTs and PDO samples at different passages. Overall, the PT-PDO copy-number discordance significantly increased over passaging (**Fig. 1E,F** and **Supplementary Fig. 3A,B**), reflecting the ongoing genomic evolution throughout PDO culture propagation. Moreover, we assessed the PT-PDO genomic discordance (**Fig. 1G**) and chromosome-arm discordance (**Supplementary Fig. 3C**) in PDO models for which data were available from multiple passages as well as from the PT of origin. We found a significant (p=0.0002; one-sample t-test), albeit modest, trend toward increased discordance throughout passaging (**Fig. 1G** and **Supplementary Fig. 3C**). These results were not associated with the tumor purity of the PT and the PDO (**Supp. Fig. 3D-G**).

Although we are not powered to detect recurrent alterations – and therefore cannot distinguish genetic drift from selection – we note that the PDO-acquired CNAs may be of functional importance, similar to what we previously described in 2D cell lines[4] and in PDXs[6,34]. Consistent with this notion, we found that cancer-related genes[35] were significantly enriched among the genes included within differential CNAs, with oncogenes such as RAD21 and CDH17 being commonly gained in PDOs and tumor suppressor genes such as *TP53, BCL2* and *CDKN2A* being commonly lost in PDOs compared to the primary tumors (**Fig. 1H**).

PDOs and PDXs are complementary preclinical models that enable the study of tumor biology and therapeutic responses in a patient-specific context, with PDXs preserving *in vivo* tumor-stroma interactions and PDOs offering a scalable, genetically faithful *in vitro* platform[14,36]. We therefore set out to assess the differences in genomic fidelity and evolution between PDO and PDX models. First, we compared the genomic CNA discordance between matched PT-PDOs and PT-PDXs originating from the same primary tumors and found that PDOs were significantly more similar to their tumors of origin. Their overall CNA discordance (**Fig. 2A**) was smaller across the three available cancer types (bladder, prostate and colorectal cancers), with a median difference of ∼8% of the genome altered in PDOs compared to ∼21% in PDXs. Similarly, the number of differentially altered chromosome-arms was significantly smaller in PDOs in comparison to PDXs (**Supplementary Fig. 4A**), with a median of 2.5 arms altered in PDOs and 6.5 arms in PDXs. We then repeated the analysis in non-matched PT-PDO and PT-PDX cohorts, which also confirmed that PDOs were more similar in their CNAs to their PTs of origin in comparison to PDXs (**Fig. 2B**), with a median genomic discordance of ∼9% in the PT-PDO cohorts compared to ∼13.5% in the PT-PDX cohorts. The median number of differentially altered chromosome-arms was also smaller in the PDOs, but only for 4 of the 6 cancer types examined (**Supplementary Fig. 4B**).

**Fig 2.**
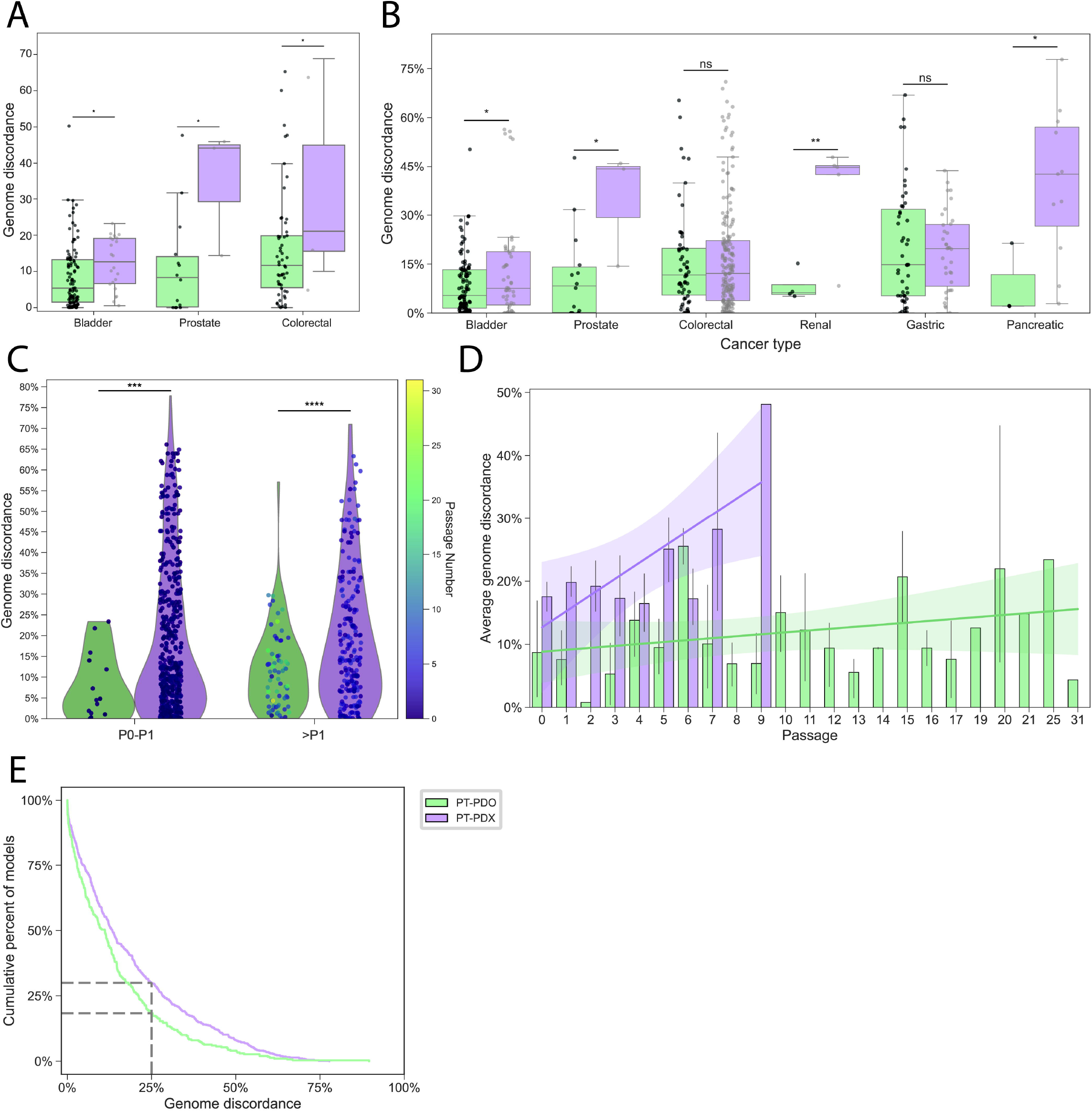
Comparison of the genetic fidelity and stability of PDOs vs. PDXs. **(A)** A cross-cohort comparison of copy-number discordance between matched PT-PDOs and PT-PDXs. The fraction of the genome differentially altered in PDOs in comparison to their PTs of origin is significantly lower than that of PDXs. Bar, median; colored rectangle, 25th to 75th percentile; whiskers, Q1–1.5*IQR to Q3+1.5*IQR. One-sided t-test; *, Bladder cancer, p-value = 0.04; Prostate cancer, p-value = 0.012; Colorectal cancer, p-value = 0.04. **(B)** A cross-cohort comparison of the CNA discordance between non-matched PT-PDOs and PT-PDXs. Across cancer types, the fraction of the genome altered is significantly lower in PDOs than in PDXs, when non-matched samples of the same tumor type are compared. Bar, median; colored rectangle, 25th to 75th percentile; whiskers, Q1–1.5*IQR to Q3+1.5*IQR. One-sided t-test; **, Renal, p-value = 0.006; *, Bladder, p-value = 0.013; Prostate, p-value = 0.012; Pancreatic, p-value = 0.023; ns., Colorectal, p-value = 0.387; Gastric, p-value = 0.618. **(C)** Comparison of the genome discordance of non-matched PT-PDOs and PT-PDXs immediately after derivation (P0-1) and in later passages (>P1, median PDO passage = 9.5, median PDX passage = 4). One-sided t-test; ****, p>1, p-value < 0.0001; ***, p0-1, p-value = 0.0001. In both passage groups the PDOs are significantly more similar to their PTs than PDXs. **(D)** A comparison of the average genome discordance between non-matched PT-PDOs and PT-PDXs across passages. PDOs present a lower discordance from the PTs across passages. One-sided Mann Whitney U test; ***, p-value = 0.0018. **(E)** A reverse estimator of a cumulative distribution function plot showing the percent of models in which over a given percentage of the genome is discordant. The PDX models were overall more genetically distant from their PTs, in comparison to PDOs. One-sided Mann Whitney U test; ***, p-value = 0.00016.

Next, we compared the discordance of PT-PDOs and PT-PDXs immediately after derivation (passage 0-1) and throughout passaging. Both at the earliest passages and after propagation, PDOs were significantly more similar to the PTs than PDXs (**Fig. 2C** and **Supplementary Fig. 4C**). Immediately after derivation (passage 0/1), the median genome discordance in PT-PDOs was ∼4.7%, in comparison to ∼11.8% (>2-fold) in PT-PDXs (**Fig. 2C**). Similar results were observed for chromosome-arm level difference: a median of ∼0.5 chromosome-arms discordant in passage 0/1 of PDOs vs. ∼2 chromosome-arms discordant in passage 0/1 of PDXs (**Supplementary Fig. 4C**). As expected, passage numbers for PDXs were on average much lower than those for PDOs (median of passage of 1 in the PDX vs. 8 in the PDOs; **Fig. 1B** and **Supplementary Fig. 4D**). Nonetheless, the genomic evolution of PDOs is slower than that of PDXs, with a smaller fraction of the genome deviating in its CN from the original PT over passaging (**Fig. 2C,D**, **Supplementary Fig. 4E**). Lastly, we calculated cumulative distribution functions for the overall CNA differences between PDO-PT and PDX-PT samples. Indeed, the PDO cohort as a whole was significantly more representative of the copy number landscape of the PTs of origin; ∼18% of the PDO models exhibited >25% genome discordance from their PTs of origin, in comparison to ∼30% of the PDX models (**Fig. 2E**).

## Discussion

In summary, we characterized the genetic CNA stability of a large cohort of PDOs generated by different labs across various cancer types and compared it to that of PDXs. Our findings revealed considerable variability in PDO similarity to their PTs of origin, even within the same cancer type, indicating that the PDO derivation protocols and/or the culture conditions are crucial for retaining the genomic stability of the models. We also found that the CN landscapes of PDOs gradually became more distant from their PTs over passaging, reflecting *in vitro* genetic evolution, consistent with the genomic evolution of all other cancer models studied to date[3–6]. These findings emphasize the importance of tracking and reporting passages while working with cancer models, and with organoids specifically. Of note, of the studies/datasets used in this work, only two directly reported the passage numbers at which the PDOs were sequenced, and ∼40% did not report clear passaging criteria (e.g., confluence or time between passages).

The *in vitro* evolution of the PDOs may have important functional consequences. First, we found that CNAs that are differential between PTs and PDOs are enriched for oncogenes and tumor suppressor genes. Second, we previously reported considerable differences in drug response between 2D cell line models that differ in ∼20% of their genetic composition[4], and between PDXs[6]. Our current findings show that on average, most PDO models stay below the 20% genome discordance threshold even at high (∼20) passages. However, individual PT-PDO discordance values may be affected by the characteristics of the primary tumor[37], its basal instability rate[14] or the culture conditions[38].

Notably, several studies previously reported stability of drug response following limited PDO passaging[39–41]. However, the effect of prolonged passaging remains to be examined more systematically. When comparing PDOs to PDXs, we found that PDOs better represented the CNA landscapes of the original tumors, and that the rate of evolution (per passage) was lower in PDOs. It’s important to note that PDO and PDX passages are not directly comparable. For example, passaging of a PDX typically involves growing a small tumor fragment to a certain size, while passaging of a PDO involves less extensive growth of cells in a well. Moreover, PDX protocols often involve passaging and sequencing of small pieces cut from the resultant tumor, whereas PDO passaging and sequencing are typically done using cells collected homogeneously from the well. Nonetheless, our findings suggest that passaging of PDOs involves less selection/drift than that of PDXs.

Together, our findings indicate that the considerable efforts invested in developing PDO derivation protocols, and culture conditions that mimic the signaling pathways to which the tumors are exposed in the human body[18,21,42], may result in smaller differences between the PT and the PDO environment, reducing the selection pressures and/or bottleneck-induced drift that lead to the genomic evolution of the models.

## Conclusions

Overall, our work provides the first comprehensive characterization of the fidelity and stability of PDOs at the CNA level, and a comparison of the two most widely used cancer-derived models. These findings are important for the application of PDOs in basic research and in translational studies.

## Methods

### Organoid Derivation

#### Human tissue

Fresh tumor specimens were obtained from patients diagnosed with malignant endometrial, breast, or bladder cancers treated at the IRCCS-Regina Elena National Cancer Institute (IRE). The study was approved by the Institutional Ethics Committee, and all participants provided written informed consent prior to sample collection and use for research purposes.

#### Establishment of organoid culture

Collected samples were maintained in MACS Tissue Storage Solution (130-100-008, Miltenyi Biotec) supplemented with 100 U/ml penicillin, 100 mg/ml streptomycin and 100 mg/ml antimycotic for a maximum of 24 hours at 4°C. Subsequently, the samples were washed twice in PBS, mechanically minced in a petri dish, and the small pieces were transferred to gentleMACS C Tubes (130-093-237, Miltenyi Biotec) or storage at -80°C tissue into the cryotubes with 1 ml of MACS Freezing solution for each cryovial (130-129-552, Miltenyi Biotec). The single-cell suspensions were obtained using Tumor Dissociation Kit (130-095-929, Miltenyi Biotec), according to the manufacturer’s instructions. The obtained cellular suspension was then filter with a 70 µm strainer (130-098-462, Miltenyi Biotec). Dissociated cell clusters were spun down at 1200 rpm for 5 min, washed once with PBS, and spun down again at 1200 rpm for 5 min. If the pellet showed a visible red color, erythrolysis was performed with Ammonium Chloride Solution (07850, STEMCELL Thecnologies) before the washing step. Dissociated cell clusters were resuspended in cold Matrigel (356231, Corning) and seeded in a prewarmed 24-well plate at density of 6×10^5^ cells per 30 µl drops. The Matrigel domes were allowed to solidify for 30 min at 37°C and 5% CO_2_ before adding 500 µL of organoid culture medium per well. The basal medium consisted of Advanced DMEM/F-12 (12634010, Gibco) supplemented with 200 mM GlutaMAX (35050038, Gibco), 1 mM HEPES (15630080, Gibco), 1.25 mM N-Acetylcysteine (A9165, Sigma Aldrich), 5 mM Nicotinamide (N0636, Sigma Aldrich), 1X N2 supplement (17502001, Invitrogen), 1X B27 supplement (17504001, Invitrogen), 250 nM A83-01 (S7692-25MG, HumanKine), 100 nM SB202190 (S7067, Sigma Aldrich), and 10 μM Y-27632 (HY-10583-10MG, HumanKine). Depending on the tumor tissue of origin, specific growth factors and supplements were added or omitted to optimize culture conditions and sustain lineage-specific signaling.

The culture medium was refreshed every 2–3 days, and organoids were passaged every 5–9 days according to their proliferation rate.

### Whole exome sequencing

DNA was extracted from frozen patient tissue and organoids using AllPrep DNA/RNA/miRNA Universal Kit (80224, QIAGEN) according to the manufacturer’s instructions. Genomic DNA was quantified using Qubit dsDNA BR Assay Kit (Invitrogen, Carlsbad, CA, USA). Quality was determined (DIN range from 1 to 10) on 4200 TapeStation using Genomic DNA screenTape assay (Agilent Technologies, Santa Clara, USA).

Pre-enrichment libraries were performed using 100 ng of DNA according to Library Preparation EF 2.0 with Enzymatic Fragmentation and the Twist Universal Adapter System (Twist Bioscience, San Francisco, CA, USA) according to the manufacturer’s instructions. Exome hybridization conducted using a Twist Comprehensive Exome kit (Twist Bioscience, CA) according to the manufacturer’s protocol. This protocol provides coverage for more than 99% of protein-coding genes. The quality of the libraries was assessed using the Agilent 4200 TapeStation system (High Sensitivity D1000 ScreenTape assay), while their quantity was measured by qPCR. The exome library was sequenced on the Illumina NovaSeq 6000 (Illumina, San Diego, CA, USA) platform with 100bp paired-end reads.

### Data assembly

Published papers in which cancer PDOs were derived and sequenced were identified using a PubMed search query containing a combination of alternative terms for “PDO” (“organoid”, “tumoroid”, “3D model”, etc.), “cancer” (“tumor”, “malignant”) and “sequencing” (“whole exome sequencing”, “WES”, “DNA sequencing”, “copy number”, “CNA”). Data for organoids which were subjected to genetic manipulation, radiation or chemical treatment prior to sequencing, or have no matched tumor sample, were removed.

Whole exome sequencing data were obtained from the European Genome-Phenome Archive (EGA) (https://ega-archive.org/), from the Database of Genotypes and Phenotypes (dbGaP) (https://dbgap.ncbi.nlm.nih.gov/home), from cBioPortal (https://www.cbioportal.org/) or directly from the corresponding authors. Data sources and accession numbers are provided in Supplementary Table 1. Normalized matrix files were downloaded, and samples were curated manually to identify the sample type and the passage number (when available). The final dataset consisted of 261 PDO samples from 183 tumors (“PDO models”) derived from 169 patients from 12 individual studies across 15 cancer types. The data was processed separately for each cohort.

### Data Processing

For each cohort, copy number alteration (CNA) data were obtained either from raw whole-exome sequencing (WES) or from previously processed segmentation tables (in original studies). When raw WES data were available, FASTQ files were aligned to the human reference genome GRCh38/GRCh37 using BWA-MEM[43] (v2.0), with lane-aware processing to maintain proper read group information. Aligned reads were coordinate-sorted, and PCR duplicates were marked using Samtools[44] (v1.10). CNA profiles were generated using CNVkit (version 0.9.10)[33] and using the reference genome build reported in the corresponding original study. The specific parameters used for the processing of each cohort are provided in the GitHub repository (https://github.com/BenDavidLab/OrganoidsCNData.git). Copy number segmentation was conducted using circular binary segmentation (CBS). The resulting CNVkit outputs, as well as segmentation tables from cohorts with pre-processed CNV data, were standardized into the. Seg format to ensure consistency across datasets. From the standardized segment files, two levels of CNA quantification were performed. Arm-level copy number values were computed by mapping genomic segments to chromosome arms and calculating length-weighted mean log_2_ ratios for each arm. In parallel, bin-level copy number values were generated by dividing each chromosome into contiguous 1Mb bins (coordinates based on the appropriate reference genome) and calculating length-weighted mean values within each bin. All processed outputs were compiled into matrices suitable for downstream statistical and comparative analyses. The full processing pipeline, including scripts and parameter specifications for each cohort, is available in the GitHub repository.

For patients with multiple tumors, each tumor was regarded as an individual model. For tumors with multiple tumor-derived PDO samples, each organoid was regarded as an individual data point in sample-level analysis, and the average of all organoid samples was used for model-level analysis.

### Computing CNA discordance between samples

CNA discordance was computed as previously reported[2,4,6]. Data were analyzed using Python V13.12.1. Copy number gains and losses were defined as |*log*2(*CN ratio*)| ≥ 0.3, respectively. For a 1□Mb bin or a chromosome-arm to be discordant between two samples, one sample must uniquely have a CN gain or loss, and additional criteria specifying the difference in log_2_(CN ratio) required between the two samples must be satisfied, to exclude borderline cases. In full, if two samples have log_2_(CN ratio) values A and B, the samples are discordant if at least one of the following conditions are met:

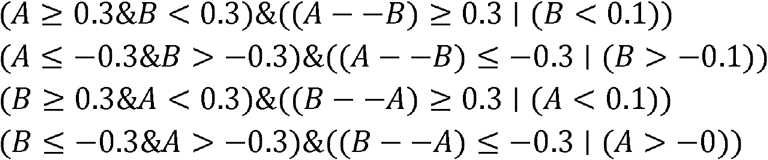

The percentage of 1□Mb bins discordant across the genome between two samples is defined as the number of discordant bins divided by the number of bins across the genome for which both samples have log_2_(CN ratio) values, multiplied by 100. The number of chromosome-arms discordant is defined as the sum of all discordant chromosome-arms between two samples.

### Purity estimation

The fraction of tumor cells in a sample was estimated from sequencing data (.bam or .fastq files) using ichorCNA, as previously reported[45].

### Computing gene discordance between samples

For each 1 Mb bin, copy number gains and losses were defined as described above. Genes were regarded as gained or lost if residing within gained or lost bins, respectively. Cancer genes were determined by the COSMIC Cancer Gene Census (https://cancer.sanger.ac.uk/cosmic/download/cosmic/v103/cancergenecensus).

The enrichment for cancer-related genes was assessed using one tailed Fisher’s exact test performed on the genes that were altered in >10% of the samples. This analysis was performed twice, to separately assess the enrichment for oncogenes within gained regions and of tumor suppressor genes within lost regions in PDOs compared to the primary tumors.

## Declarations

## Acknowledgments

We thank the individuals and research groups who made the data used in this study available. Specifically, we thank Johnathan Yeung, Suet Yi Leung, Marianna Kruithof De Julio, Rebekah Engel, Helen Abud and Tomohiko Ishiwaka for providing direct access to the data from their studies[22–25,27,38]. We also thank the authors who shared their studies through the European Genome-phenome Archive (EGA)[28,29]. Finally, we thank the authors who made their data publicly available through cBioPortal or directly from the paper [30–32].

## Author Contributions Statement

U.B.-D. conceived the study and supervised the project. L.R. led the data collection, performed the analyses, and generated the plots and the table. H.R., T.B.Y. and R.S. processed the sample sequencing data and contributed to data collection and analysis. A.Z. participated in data collection. J.O., S.D. and G.B. created the new PDOs reported in this work. L.R. and U.B.-D. wrote the paper, with inputs from all co-authors.

## Data availability statement

All data supporting the findings of this study are available within the paper and its Supplementary Information. All custom code is available at https://github.com/BenDavidLab/OrganoidsCNData.git.

## Supplementary Figure Legends

**Supplementary Fig. 1.Further information on the PDOs analyzed in this study.**

**(A)** Cancer type composition of the PDO models included in this study. Each model represents one or more samples derived from the same PT. **(B)** Distribution of all matched samples used in this study. Matched samples are derived from the same primary tumor and represent PTs and their derived PDOs (PT-PDO), PTs and their derived PDOs and PDXs (PT-PDO-PDX) or PTs and their derived PDOs across several passages.

**Supplementary Fig. 2. PT-PDO CNA discordance across cohorts.**

**(A**,**B)** Precent of the genome that is discordant **(A)**, or the number of chromosome-arm CNAs that are discordant **(B)**, between PDOs and their PTs of origin across cancer types, shown for PDO models (rather than for individual samples). Each data point is the average of all PDO samples derived from a single tumor. The analysis includes only cohorts with at least 3 models. Bar, median; colored rectangle, 25th to 75th percentile; whiskers, Q1–1.5*IQR to Q3+1.5*IQR. **(C**,**D)** Percent of the genome that is discordant **(C)**, or the number of chromosome-arm CNAs that are discordant **(D)**, between PDOs and their PTs of origin across the different cohorts included in this study. Boxplot color indicates cancer type. Bar, median; colored rectangle, 25th to 75th percentile; whiskers, Q1–1.5*IQR to Q3+1.5*IQR.

**Supplementary figure 3.PDO genomic divergence throughout passaging.**

**(A**,**B)** The percentage of genome that is discordant **(A)** or the number of chromosome-arm CNAs that are discordant **(B)** between the earliest and latest matched organoid samples derived from the same tumors. Across most samples, the discordance increases over passaging at both the FGA-level level and the chromosome-arm level. **(C)** The cumulative PT-PDO chromosome-arm discordance in a longitudinal series of matched PDOs. Each segment is colored by the change in discordance between the two consecutive organoids. Most passaging steps lead to a change in discordance smaller than 1 chromosome arm. One-sided one-sample t-test; p-value < 0.0001. **(D)** Comparison of PT and PDO purity. PDO samples are significantly purer than tumor samples. Two-sided t-test; ****, p-value < 0.0001. **(E)** Correlation between PT-PDO genome discordance and PDO sample purity. Spearman’s correlation; Rho = 0.1247, p-value = 0.4893 **(F)** Correlation between PT-PDO genome discordance and tumor sample purity. Spearman’s correlation; Rho = 0.1235, p-value = 0.4933 (g) Correlation between PT-PDO genome discordance and the difference between PDO and tumor sample purity. Spearman’s correlation; Rho = 0.0276, p-value = 0.8789.

**Supplementary Fig. 4. Comparison of chromosome-arm CNA fidelity and stability between PDOs and PDXs.**

**(A)** A cross-cohort comparison of chromosome-arm CNA discordance between matched PT-PDOs and PT-PDXs. PDOs display a significantly lower percentage of the genome altered. One-sided t-test; ns, Bladder, p-value = 0.078; Prostate, p-value = 0.052; Colorectal, p-value = 0.112. **(B)** A cross-cohort comparison of the CNA discordance between non-matched PT-PDOs and PT-PDXs. Across cancer types, the number of chromosome-arm CNAs altered are lower in PDOs than in PDXs, when non-matched samples of the same tumor type are compared. Bar, median; colored rectangle, 25th to 75th percentile; whiskers, Q1–1.5*IQR to Q3+1.5*IQR. One-sided t-test; *, Renal, p-value = 0.01; ns, Bladder, p-value = 0.238; Prostate, p-value = 0.052; Colorectal, p-value = 0.81; Gastric, p-value = 0.996; Pancreatic, p-value = 0.065. **(C)** Comparison of the chromosome-arm discordance of non-matched PT-PDOs and PT-PDXs immediately after derivation (P0-1) and in later passages (>P1, median PDO passage = 9.5, median PDX passage = 4). One-sided t-test; ns, p-value = 0.1439; *, p-value = 0.0301. In both passage groups the PDOs are significantly less discordant from their PTs than PDXs. **(D)** Distribution of PDX passages across samples. **(E)** A comparison of the average number of chromosome-arms that is discordant between non-matched PT-PDOs and PT-PDXs across passages. While PDOs seem to exhibit a lower number of arms discordant from the PTs across passages, the trend is not significant. One-sided Mann Whitney U test; ***, p-value = 0.529.

## Table Legends

**Supplementary Table 1. Description of organoid cohorts and samples used in this study.** For each study, shown are the cancer type(s), the numbers of patients/models/samples, the source of the data and the publication reporting it, whether passage numbers were reported in the study, and – when available – the passage numbers and passaging method used in the study.

