## Supplementary figures and images for "Genomic fidelity and evolution of cancer patient-derived organoids"

### Supplementary figure 1

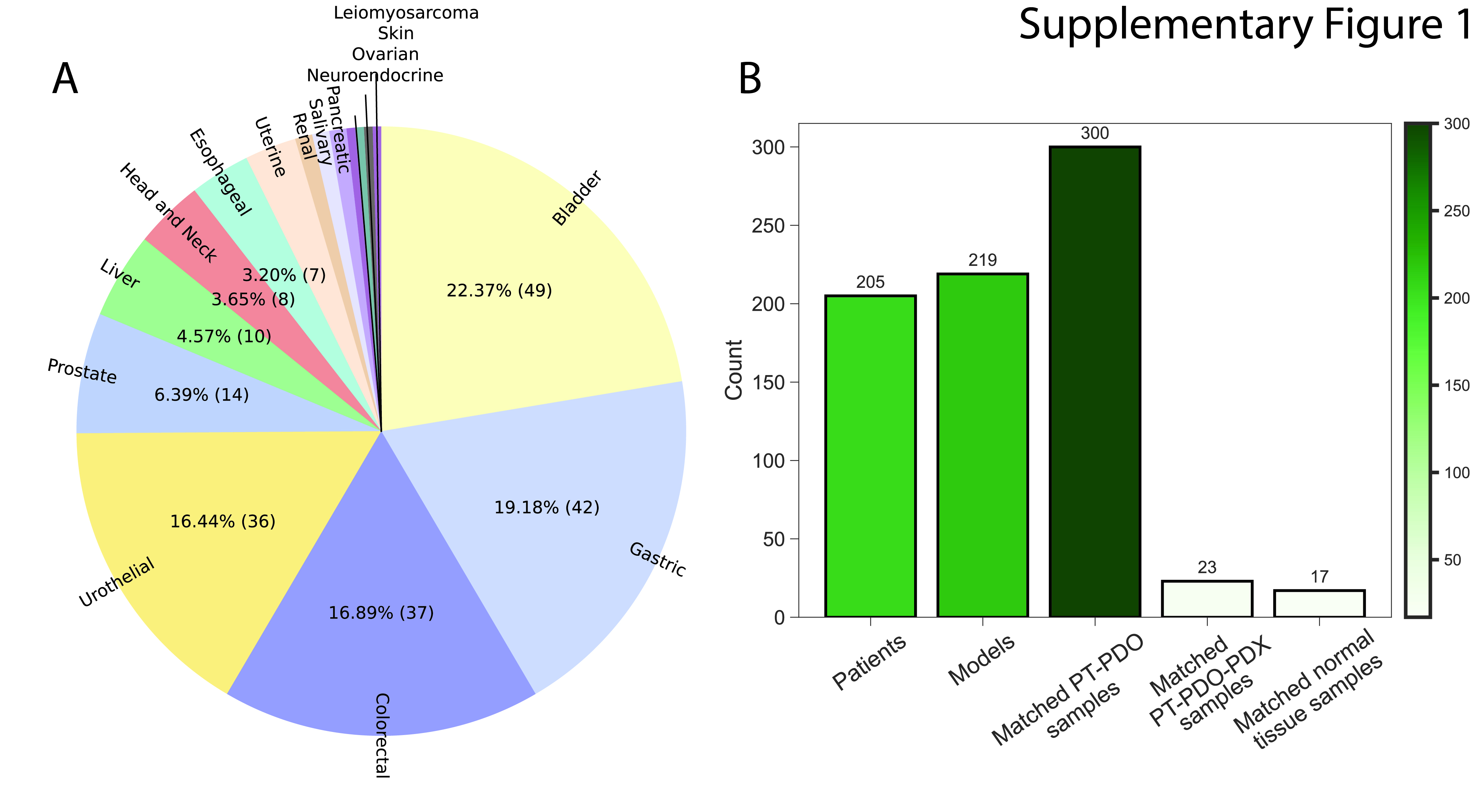

### Supplementary figure 2

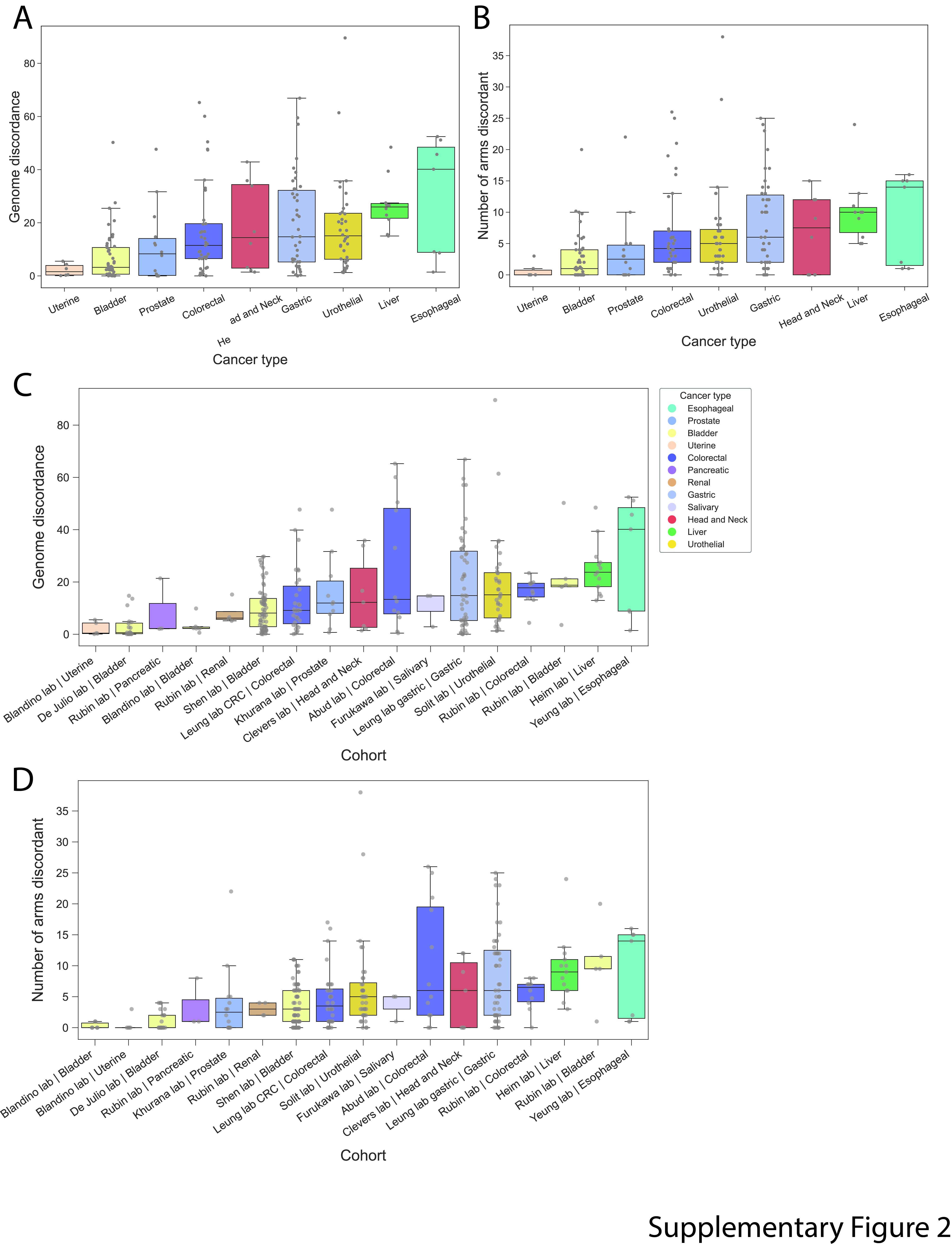

### Supplementary figure 3

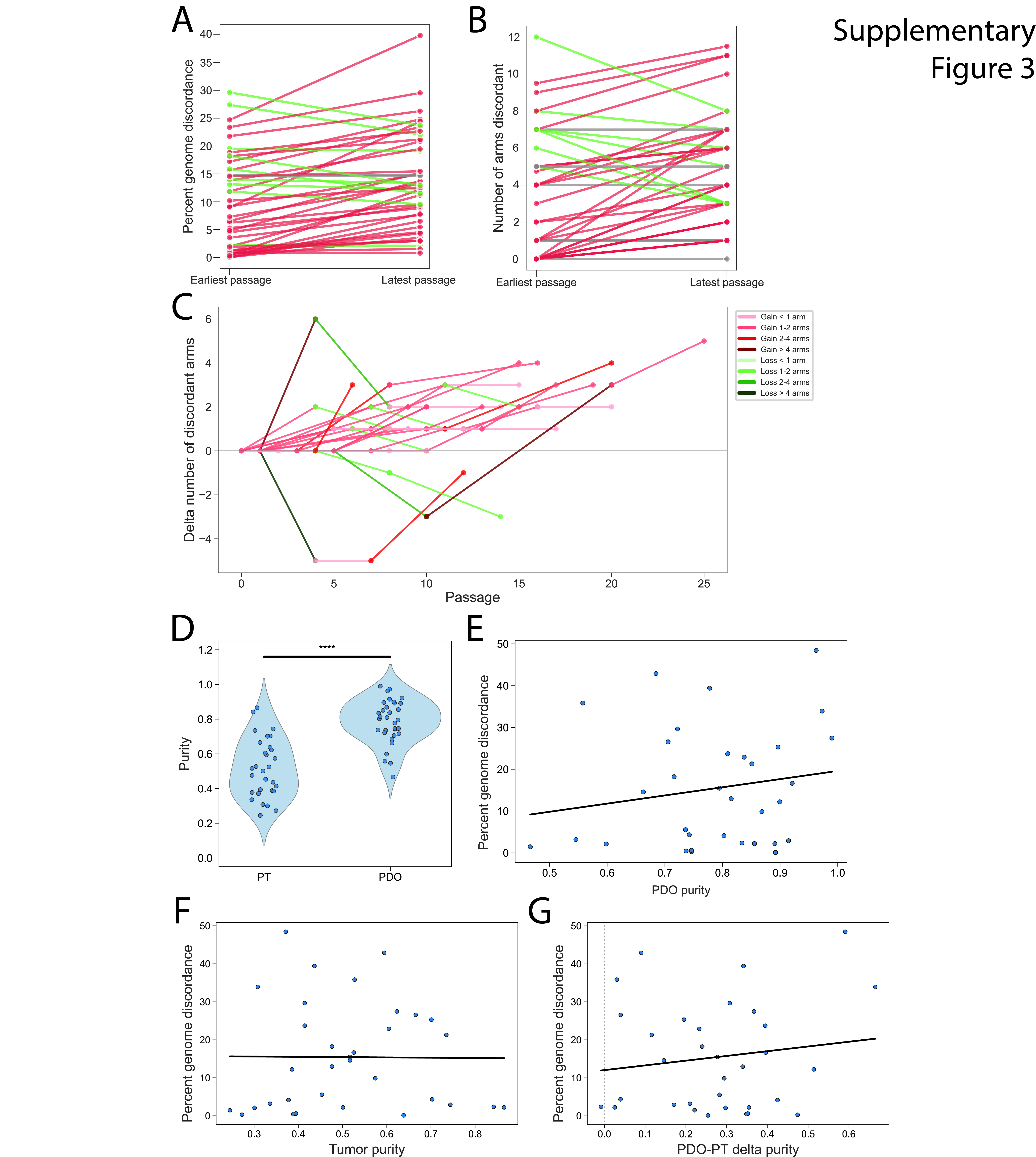

### Supplementary figure 4

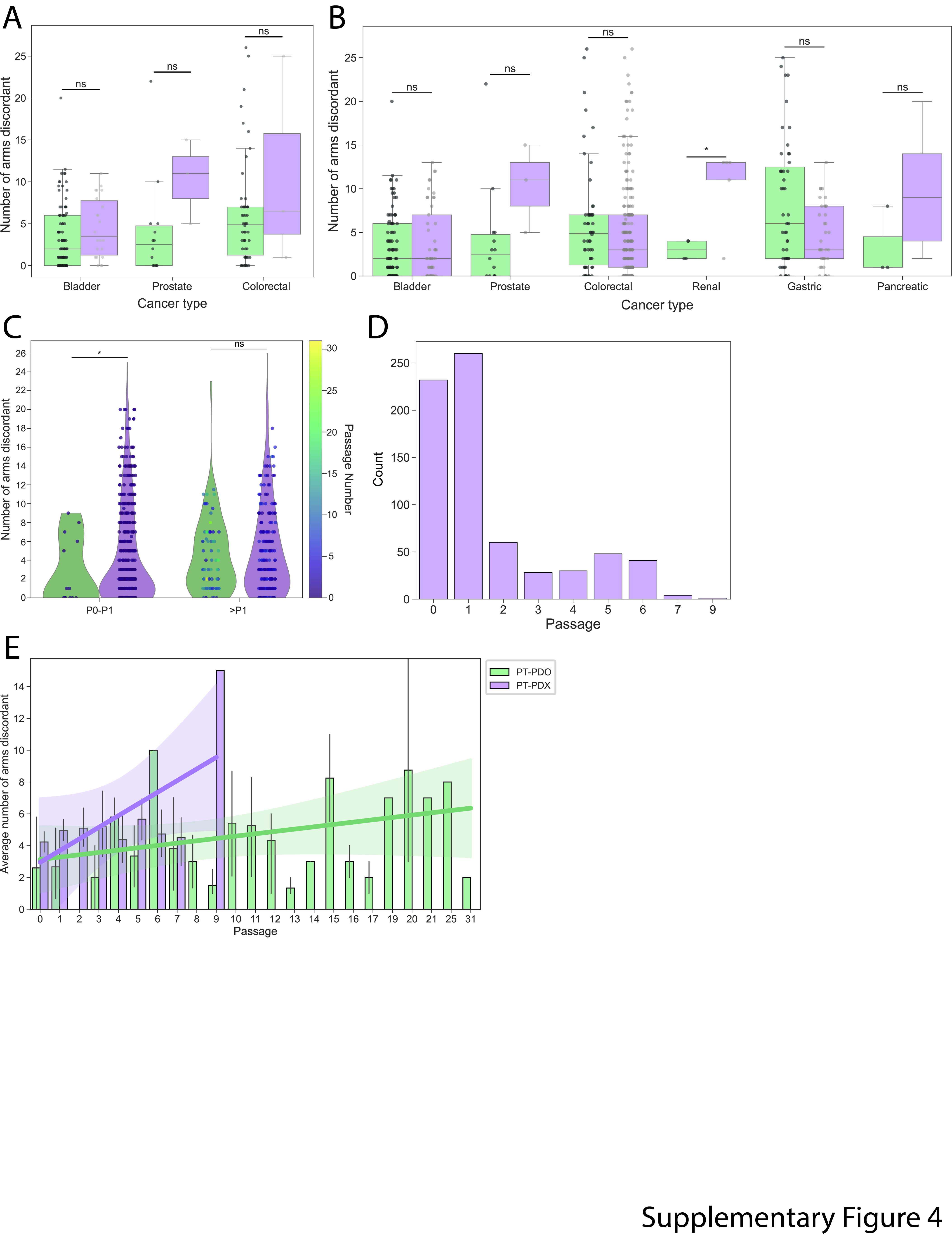
